# Assessing the efficiency and heritability of blocked tree breeding trials

**DOI:** 10.1101/2024.06.01.596996

**Authors:** Hans-Peter Piepho, Emlyn Williams, Maryna Prus

**Author notes:** **Correspondence** Hans-Peter Piepho, Biostatistics Unit, Institute of Crop Science, University of Hohenheim, 70593 Stuttgart, Germany.

## Abstract

Progeny trials in tree breeding are often laid out using blocked experimental designs, in which families are randomly assigned to plots and several trees are planted per plot. Such designs are optimized for the assessment of family effects. However, tree breeders are primarily interested in assessing breeding values of individual trees. This paper considers the assessment of heritability at both the family and tree levels. We assess heritability based on pairwise comparisons among individual trees. The approach shows that there is considerable heterogeneity in pairwise heritabilities, primarily due to the differences in both genetic as well as error variances among within- and between-family comparisons. Our results further show that efficient blocking positively affects all types of comparison except those among trees within the same plot.

## Introduction

Progeny trials, commonly employed by tree breeders, are usually laid out as randomized complete block designs (RCBD) or resolvable incomplete block designs (ICBD), where each plot has several trees, originating from the same family. The design has families (either open-pollinated, or control-pollinated) as the treatment factor whose levels are allocated to plots, meaning that the design is optimized for the estimation of family effects. However, breeders are interested not only in the family means, but also in the breeding values of individual trees, because there is also genetic variance to be exploited within families and selection is commonly exercised on individual trees (forward selection, or sometimes backward selection; White et al. 2007).

Derivations of the formulas for heritability used in tree breeding (for example Namkoong et al. 1988; White et al. 2007; Williams et al. 2024) are based on RCBD progeny trials, generally employed prior to the widespread adoption of ICBDs over the last 2-3 decades. The present paper is concerned with an assessment of the efficiency of blocking when estimating breeding values of individual trees. Furthermore, we propose methods to estimate heritability on an individual-tree basis. One real example from a progeny trial is considered for illustration. Our initial focus is on experiments laid out according to a RCBD, where we assume that the same number of trees is available in each plot. This simple setting serves to introduce our basic concepts and to explore key properties of our proposed approaches using explicit scalar equations available in this case. We consider two different approaches for assessing heritability, one based on the variance explained by genetic effects and one based on the squared correlation between genetic values and their best linear unbiased predictors (BLUP). Subsequently, we consider the more common situation that there is some mortality over the course of the experiment, meaning that the number of trees varies among plots. In this case, we only employ the BLUP-based approach, which we recommend for routine use. Finally, we explore more complex systems of blocking, including alpha designs and resolvable row-column designs (Williams et al. 2024), also focusing on BLUP. The salient feature of our proposed approaches is the focus on pairwise differences (Schmidt et al. 2019).

## Material and Methods

### Two equivalent models for trials laid out as RCBD

For an experiment laid out as an RCBD, we may use the model

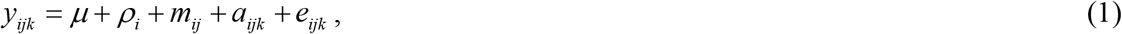

where *y*_*ijk*_ is the response of the *k*-th tree of the *j*-th family in the *i*-th replicate (*i* = 1,…*I*; *j* = 1,…, *J*; *k* = 1,…, *K*), *μ* is an overall intercept, *ρ*_*i*_ is the fixed effect of the *i*-th replicate, 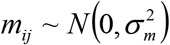 is the random effect of the *j*-th plot in the *i*-th replicate, *a*_*ijk*_ is the breeding value (additive genetic effect) of the *ijk*-th tree, and 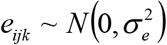 is a residual associated with *y*_*ijk*_, comprising both residual genetic effects and non-genetic errors.

The breeding values are assumed to be multivariate normal with a correlation of *r* within families and no correlation between families (Falconer and Mackay 1996, p.233). Specifically, *r* is assumed to be ¼ for half-sib families, and ½ for full sib families. Our focus is on open-pollinated progeny trials, where *r* is increased from ¼ up to ⅓ or more in order to account for levels of selfing, inbreeding among relatives and correlated paternity; this will vary according to species breeding biology and particular circumstances. Here, we will use the value *r* = 0.3 (Williams et al. 2024, p.96). The value of *r* can be fixed in the analysis, meaning that only the variance of 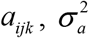, needs to be estimated. This model implies a compound symmetry variance-covariance structure for the within-family correlation, and this can be exploited in fitting the model with a mixed model package. The advantage of model (1) is that the BLUPs of *a*_*ijk*_ can be obtained directly from the mixed model equations.

An alternative but equivalent form of the model is given by

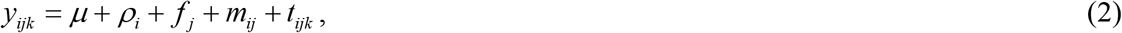

where 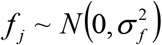 is a random family effect, and 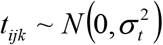 is a random tree effect. The equivalence relations of the effects between models (1) and (2) are *a*_*ijk*_ = *fj* + *b*_*ijk*_ and *t*_*ijk*_ = *b*_*ijk*_ + *e*_*ijk*_. For the variances, we have the equivalence relations 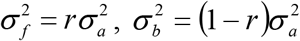, and 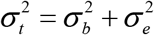.

Under model (2) we are confounding additive genetic effects *b*_*ijk*_, and non-additive genetic effects as well as non-genetic error effects *e*_*ijk*_ in the tree effect *t*_*ijk*_. Obviously, we must have 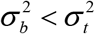. Using the equivalence relations for the variances as stated above, we obtain the condition

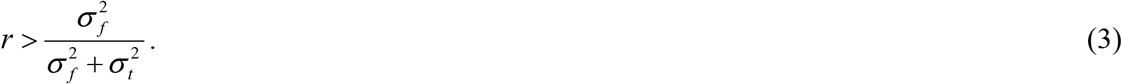

This suggests that one should initially fit model (2) to check if condition (3) is met. If this is the case, one may proceed to fit model (1).

### Pairwise heritabilities for RCBD

#### Family means basis – complete data

Heritability may be assessed on a family-mean basis or on an individual-tree basis. We first consider family-mean based heritability. There are two different approaches to heritability assessment that can be pursued here. One option is to consider the best linear unbiased estimator (BLUE) of the family mean, given by 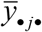 in the case of an RCBD with complete data, and assess the proportion of the phenotypic variance among these means that is explained by the family effect. Note that we define complete data as all trees being present in each plot. For an RCBD with complete data, the family-mean heritability is given by

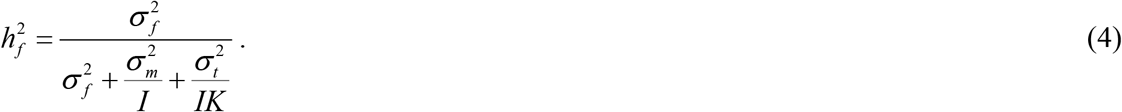

Note that we may also consider (4) as the slope for the regression of the family effect *f*_*j*_on the family mean 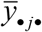 Family-mean heritability is used to predict gain obtained by selecting a proportion of the families under test, without breeding from them. In this sense it is the same as heritability for crop varieties, where variety means are computed across replicates per trial (Piepho and Möhring 2007).

The second option for family-means is to define heritability as the squared correlation between the true family effect *f*_*j*_ and its BLUP. This BLUP-based approach is conceptually more closely connected to the response to selection (Falconer and Mackay 1996, p. 232), which is a key concern for breeders. For the complete data case considered here, this BLUP-based definition leads to the same equation as given in (4) for the BLUE-based approach. This can be shown based on the fact that the BLUP of *f*_*j*_ is 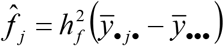, and working out the squared correlation between *f*_*j*_ and 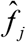.

Heritability is most often introduced on the basis of an individual genotype (e.g., a tree) or an individual family. An alternative view, which we prefer, is to consider pairs of genotypes or families (Schmidt et al. 2019). Clearly, the difference among two family effects determines which families should be selected. Hence, under the BLUE-based approach (Piepho and Möhring 2007), we may consider the difference 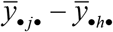 for two families *j* and *h* and derive their variance. The genetic variance of this difference is 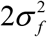 and its phenotypic variance is 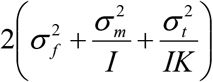. The ratio of these two variances is equivalent to equation (4). Defining the semivariance as half the variance of a difference (Feldmann et al. 2022), we may also say that the heritability in (4) is the ratio of the genetic over the phenotypic semivariance. The same result is obtained for the BLUP-based approach if applied to family differences rather than family means (Cullis et al. 2006).

Our focus on mean differences rather than means provides no novel measure of heritability in this case, because the variance of a difference among family means is the same for each pair of families. It does, however, provide a very useful way to define heritability in more complex settings (Schmidt et al. 2019), where the variance of a difference is not constant across pairs of families, as is the case when incomplete blocks are used and when the number of trees varies between plots.

#### Individual tree basis – complete data

Let us now turn to heritability at the level of individual trees. It is worth stressing at this point, that an RCBD with multiple-tree plots is optimized for the comparison among family means, but it is not explicitly optimized for the comparison of individual trees. An advantage of multiple-tree plots is better robustness than single-tree plots to tree mortality. A further aspect is a potential reduction in average inter-tree competition, relative to a single-tree plot design, because trees in a family line plot have either one, or two adjacent trees belonging to the same family, whereas in a single-tree plot design all immediate neighbours are from a different family and hence differences in height are expected to be larger. When all trees of the same family in a replicate are planted in the same plot, it is also clear that blocking helps improving the efficiency of family mean estimates because the plot error variance 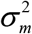, which affects the denominator in (4), will be reduced when blocking by replicates is effective. The impact of blocking on the efficiency of tree comparisons is less obvious. Again, it is useful to consider pairwise differences. As we will see, the pairwise variances of differences vary among pairs of trees.

There is only a single observation per tree, and hence, a BLUE for individual tree effects is not available. Instead, we may consider an approach that works with the observed data *y*_*ijk*_. However, when comparing trees in different replicates, we cannot use the observed data directly due to the presence of replicate effects. Instead, we may select the best trees based on replicate-corrected data 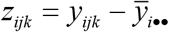, which sweep out the replicate effect (de Hoog et al. 1990). While this approach is seldom used by tree breeders, it does provide a useful framework for defining pairwise heritabilities at the tree level. Note that for an RCBD with complete data replicates are orthogonal to families, and hence a two-stage process of sweeping replicates is the same as including replicate effects in a one-stage model (Williams 1986).

There are four different types of pairwise tree comparisons:

a. Two trees in the same family and plot: There are *IJK* (*K* −1) 2 such pairs.
b. Two trees in the same family but in different replicates: There are *I* (*I* −1)*JK* ^2^ 2 such pairs.
c. Two trees in different families but the same replicate: There are *IJ* (*J* −1)*K* ^2^ 2 such pairs.
d. Two trees in different families and replicates: There are *I* (*I* −1)*J* (*J* −1)*K* ^2^ 2 such pairs.

Table 1 lists the pairwise genetic and error semivariance of a difference for these four types of comparison as derived from model (2), where the semivariance is defined as half the variance of a difference of two corrected values *z*_*ijk*_ (Piepho 2023). The phenotypic semivariance is the sum of the genetic and error semivariances. The ratio of the genetic over the phenotypic semivariance is the pairwise heritability for the replicate-corrected data. It is clear that each type of comparison has a different pairwise heritability. As pointed out before, if blocking by replicates is effective, the plot error variance 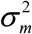 will be reduced. Hence, effective blocking potentially has a positive effect on comparisons (b) to (d), but not on comparison (a), which is a within-plot comparison. However, for comparisons (b) and (d), the adjustment for replicate effects may adversely affect pairwise variances, so the net effect of blocking will depend on the values of the variance components. Note that in comparisons (b) and (d), the genetic variance component 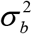 appears in both the genetic variance of the pairwise comparison of interest and the corresponding residual. This is because apart from the two genetic effects *b*_*ijk*_ of the comparison of interest, there are effects *b*_*ijk*_ of other trees in the same two replicates, and these must be regarded as residual noise. It must also be acknowledged that the replicate-corrected value *z*_*ijk*_ entail a slight bias with respect to the random effect *b*_*ijk*_, because the correction leads to a coefficient of (*KJ* − 1)/(*KJ*) for this effect due to the subtraction of the replicate mean 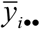. The bias will be small, however, when the product *KJ* is large. This bias-causing term drops out for pairwise comparisons (a) and (c) involving trees in the same replicate, but it persists for comparisons (b) and (d), which also shows in the coefficients for 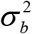 in the genetic variance in Table 1.

**Table 1:** Pairwise semivariances (genetic, error and phenotypic) for four types of comparison (a, b, c and d) based on corrected data 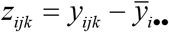 for RCBD with complete data under model (2) using *a* 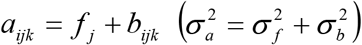 and 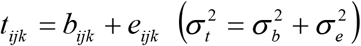 in relation to model (1). The phenotypic semivariance is the sum of the genetic and error semivariances. The ratio of the genetic over the phenotypic semivariance is the pairwise heritability for the comparison.

| Type of pairwise comparison | Semivariance |  |  |
| --- | --- | --- | --- |
|  | Genetic | Error | Phenotypic |
| (a) Two trees in the same family and plot | $\sigma_b^2$ | $\sigma_e^2$ | $\sigma_t^2$ |
| (b) Two trees in the same family but in different replicates | $\frac{(JK-1)^2}{J^2 K^2} \sigma_b^2$ | $\frac{JK-1}{J^2 K^2} \sigma_b^2 + \frac{J-1}{J} \sigma_m^2 + \frac{JK-1}{JK} \sigma_e^2$ | $\frac{J-1}{J} \sigma_m^2 + \frac{JK-1}{JK} \sigma_t^2$ |
| (c) Two trees in different families but the same replicate | $\sigma_f^2 + \sigma_b^2$ | $\sigma_m^2 + \sigma_e^2$ | $\sigma_f^2 + \sigma_m^2 + \sigma_t^2$ |
| (d) Two trees in different families and replicates | $\sigma_f^2 + \frac{(JK-1)^2}{J^2 K^2} \sigma_b^2$ | $\frac{JK-1}{J^2 K^2} \sigma_b^2 + \frac{J-1}{J} \sigma_m^2 + \frac{JK-1}{JK} \sigma_e^2$ | $\sigma_f^2 + \frac{J-1}{J} \sigma_m^2 + \frac{JK-1}{JK} \sigma_t^2$ |

The pairwise heritability for comparison (c), 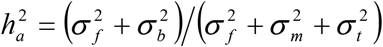, coincides with eq. (6.6) in Williams et al. (2024), 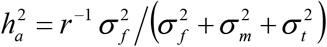 when using the equivalence relations between models (1) and (2). The pairwise semivariances for comparisons (b) and (d) look more complex because different replicates are involved, meaning that the replicate mean 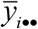 remains in the comparison. By contrast, for comparisons (a) and (c), we obtain 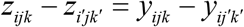, which leads to simpler expressions for the semivariance. In other words, comparison (c) also applies for the original data and thus provides some support for the commonly used expression for individual tree heritability.

Next, consider the BLUP-based pairwise heritabilities. These may be computed from the genetic semivariances and the corresponding error semivariances of BLUPs for the genetic effect *a*_*ijk*_ shown in Table 2, obtained from the inverse of the coefficient matrix of the mixed model equations (see next section). One minus the ratio of the error semivariance over the genetic semivariance is the pairwise heritability for the comparison (Schmidt et al. 2019). It can be shown from the results in Table 2 that the heritability for comparison (a) equals 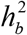, which is identical to the heritability for that comparison in Table 1. For the other comparisons, however, the heritabilities do not agree between the two tables.

**Table 2:** Error semivariances of BLUPs for genetic effect *a*_*ijk*_ and genetic semivariance for four types of comparison (a, b, c and d) for RCBD with complete data under models (1) and (2). One minus the ratio of the error semivariance over the genetic semivariance is the pairwise heritability for the comparison.

| Type of pairwise comparison | Error semivariances of BLUPs for genetic effect $a_{ijk}$ | Genetic semivariance |
| --- | --- | --- |
| (a) Two trees in the same family and plot | $\frac{1}{c_1}$ | $\sigma_b^2$ |
| (b) Two trees in the same family but in different replicates | $\frac{1}{c_1} \left( 1 + \frac{d_2}{K} + \frac{d_4}{KJ} \right)$ | $\sigma_b^2$ |
| (c) Two trees in different families but the same replicate | $\frac{1}{c_1} \left( 1 + \frac{d_2}{K} + \frac{d_3}{KI} \right)$ | $\sigma_a^2$ |
| (d) Two trees in different families and replicates | $\frac{1}{c_1} \left( 1 + \frac{d_2}{K} + \frac{d_3}{KI} + \frac{d_4}{KJ} \right)$ | $\sigma_a^2$ |

The BLUP of *a*_*ijk*_ = *f*_*j*_ + *b*_*ijk*_ can be written as the sum of the BLUP of *f*_*j*_ and the BLUP of *b*_*ijk*_. It may be shown (see Appendix S1) that for a balanced RCBD

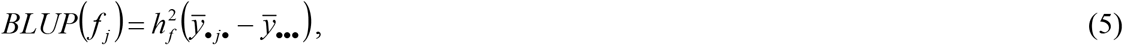

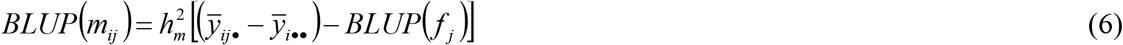

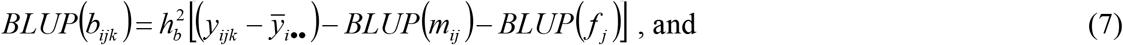

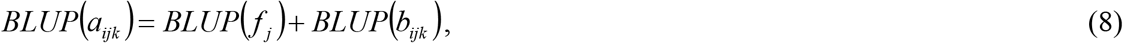

where 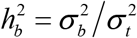 and 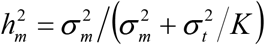. Note that (8) is essentially Lush’s (1947a,b) index for combined within and between family selection (also see Falconer and Mackay 1996, p.235), with additional terms in (6) that account for experimental design effects. From the explicit equations (5) and (6) for BLUP of *f*_*j*_ and *b*_*ijk*_, we can make some conjectures about possible gains from blocking for the BLUP of *a*_*ijk*_ = *f* _*j*_ + *b*_*ijk*_. To explore this, note again that blocking can potentially reduce the plot variance 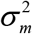, but will not affect any other variance.

i. It is clear that the BLUP of *f*_*j*_ and their pairwise contrasts will benefit from blocking, because families are the treatments that are randomly allocated to plots according to an RCBD.
ii. For the contrast of the BLUP of *b*_*ijk*_ for two trees, we need to distinguish the four cases depending on whether or not the two trees are in the same family and on whether or not the two trees are in the same block.
  a. Two trees in the same family and plot: It is clear that there is no benefit from blocking.
  b. Two trees in the same family but in different replicates: The adjustments in (7) involving the BLUPs of *m*_*ij*_ and *f* benefit from a reduction of 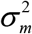, but the pairwise contrast may be adversely affected by the adjustment for replicate effects. The net effect will depend on the magnitude of the reduction of 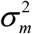.
  c. Two trees in different families but the same replicate: The replicate effect drops out in a pairwise contrast. Hence, we may expect a gain from blocking.
  d. Two trees in different families and replicates: See (b).

The error and genetic semivariances for the BLUP of *a*_*ijk*_ for the four types of comparison are given in Table 2 (see Appendix S2 for a derivation of the error semivariances).

Based on the results in Table 2, we make the following observations:

i. We have 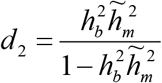 in comparisons (b), (c) and (d). This is reduced by reducing 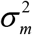, implying a benefit from efficient blocking in these comparisons.
ii. We have 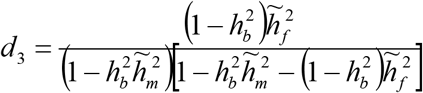 in comparisons (c) and (d), which is reduced by reducing 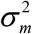, implying a benefit from efficient blocking.
iii. With 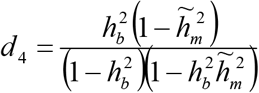 and 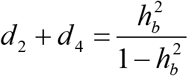, we find that 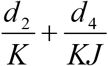 is monotonically increasing in 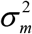, implying a further benefit from efficient blocking in comparisons (b) and (d).
iv. The pairwise heritability for comparison (b) will generally be smaller than that for comparison (a).
v. The pairwise heritability for comparison (d) will generally be smaller than that for comparison (c).

The pairwise heritabilities may be averaged across pairs to obtain an overall measure of heritability (Schmidt et al. 2019). Alternatively, one may first average the semivariances in the numerator and denominator of the ratio involved and then compute the ratio (Feldmann et al. 2022). A general method for this based on mixed model output will be described in the next section.

### A general method to compute pairwise heritabities

The use of replicate-corrected data *z*_*ijk*_ for selection could be extended to other designs with more complex blocking, such as resolvable row-column designs. All that is required are estimates of the relevant block effects. These estimates would then be subtracted from the observed data to obtain the corrected data. Assessing heritabilities based on the replicate-corrected data is possible in principle, but will not be considered here because we think the BLUP-based approach is generally preferable due to its better efficiency, as will be illustrated in the next section.

The BLUP-based pairwise heritability may be defined as (Schmidt et al. 2019)

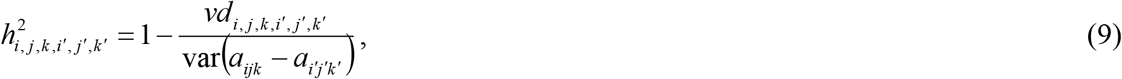

where 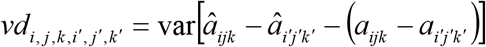 is the error variance of the difference of the two BLUPs for *a*_*ijk*_ and *a*_*i′j′k′*_. If var (*a*_*ijk*_ − *a*_*i′j′k′*_) were constant, we could average this across all pairs (Cullis et al. 2006). With variable var(*a*_*ijk*_ *− a*_*i′j′k′*_), we can make an approximation an average both var (*a*_*ijk*_ *− a* _*i′j′k′*_) and *νd*_*i,j,k,i′,j′,k′*_, leading to the average heritability

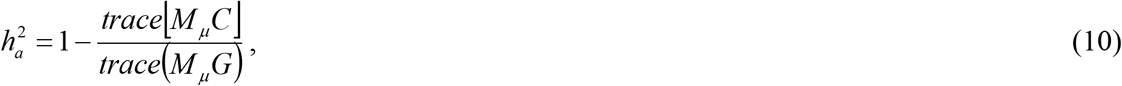

where *M* _*μ*_ = *I*_*n*_ − *K*_*n*_ is a mean-centering matrix with *I*_*n*_ the *n*-dimensional identity matrix,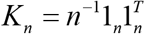 is the projection matrix (James and Williams 2024) for the grand mean, *n* = *IJK*, *C* = var(*â* − *a*), *G* = var(*a*), *a* is the vector of the random effects *a*_*ijk*_, and *â* its BLUP. The matrix *C* is obtained from the inverse of the coefficient matrix of the mixed model equations.

### An *Acacia mangium* progeny trial, South Sumatra

This trial was planted in 1995, by PT. Musi Huta Persada. It tested *J* = 56 open-pollinated families collected from the extensive natural occurrences of the species around the settlement of Wipim, in Western Province, Papua New Guinea. The trial was part of a larger breeding programme for *A. mangium* implemented by the company with technical advice from Dr E. B. Hardiyanto of Gada Madja University, which established several sub-lined breeding populations each representing a different natural provenance.

The trial had *I* = 12 replicates in a 6 × 2 arrangement and each replicate comprised 8 × 7 plots of *K* = 4 trees at 4m × 2m spacing. We have concentrated on one year diameter at breast height (dbh1) which was measured just prior to selective thinning of the trial. Survival was at 90% but one of the families (no. 57) which had a shortfall of plantable seedlings at trial establishment was padded out with similarly performing fillers in six of the 12 replicates. Hence interpretation of results for this family would be more difficult but will have no impact on our study. We will refer to the dbh1 data from the trial (i.e. with missing trees) as incomplete trial data.

### Simulated complete data

We will first use this example to exemplify results for an RCBD with complete data. To this end, we initially did not fit row and column effects. We simulated complete data from the model fitted to the incomplete data, adding random effects for rows and columns nested within replicates to represent the original randomization layout. This was done by computing replicate means from the dbh1 data and adding on random normal deviates for effects *f*_*j*_, *m*_*ij*_ and *t*_*ijk*_ in model (2) as well as random effects for rows and columns that had the same variances as those estimated from the incomplete data (Table 3).

**Table 3:** REML estimates of variance components for diameter at breast height at one year (dbh1) using simulated complete and incomplete *Acacia mangium* data (Example 1).

| Variance parameter | Simulated complete data |  | Incomplete trial data |  |
| --- | --- | --- | --- | --- |
|  | Estimate | s.e. | Estimate | s.e. |
| $\sigma_m^2$ | 0.1684 | 0.02986 | 0.1841 | 0.0337 |
| $\sigma_a^2$ | 0.3507 | 0.09301 | 0.3672 | 0.0988 |
| $\sigma_e^2$ | 1.0358 | 0.07659 | 1.0292 | 0.0816 |
| $\sigma_f^2$ | 0.1053 | 0.02795 | 0.1102 | 0.0296 |
| $\sigma_t^2$ | 1.2813 | 0.04036 | 1.2863 | 0.0434 |

The variance parameter estimates based on models (1) and (2) are reported in Table 3. For model (2) we find 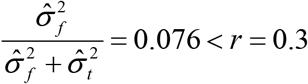, so fitting model (1) is feasible. The heritability on a family-mean basis as per eq. (4) for the simulated complete data equals 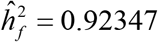. The tree-level pairwise heritabilities based on Tables 1 and 2 are shown in Table 4. For comparison (a), the heritability based on replicate-corrected values agrees with the heritability based on BLUP, equalling 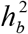 as expected [Note that BLUP is computed from the observed data *y*_*ijk*_ directly, so no correction for replicate means is needed; the correction is implicit in the BLUP equations (5) to (8)]. The small discrepancy is due to small numerical differences between the fits of models (1) and (2). For the other comparisons, there are more pronounced differences. The largest discrepancy occurs for the between-family comparisons (c) and (d). This is mainly due to the fact that BLUP borrows strength from observations in the same family, which improves the accuracy of between-family comparisons compared to the use of replicate-corrected data, which clearly do not borrow strength. We reiterate that pairwise heritability for comparison (c) in Table 1 for replicate-corrected data *z*_*ijk*_ is the same as the heritability given in eq. (6.6) in Williams et al. (2024). As pairwise heritabilities for (c) and (d) are roughly equal numerically (Table 4) and account for the majority of the pairwise comparisons, the average heritability across all pairs is very similar to that from eq. (6.6) in Williams et al. (2024).

**Table 4:**
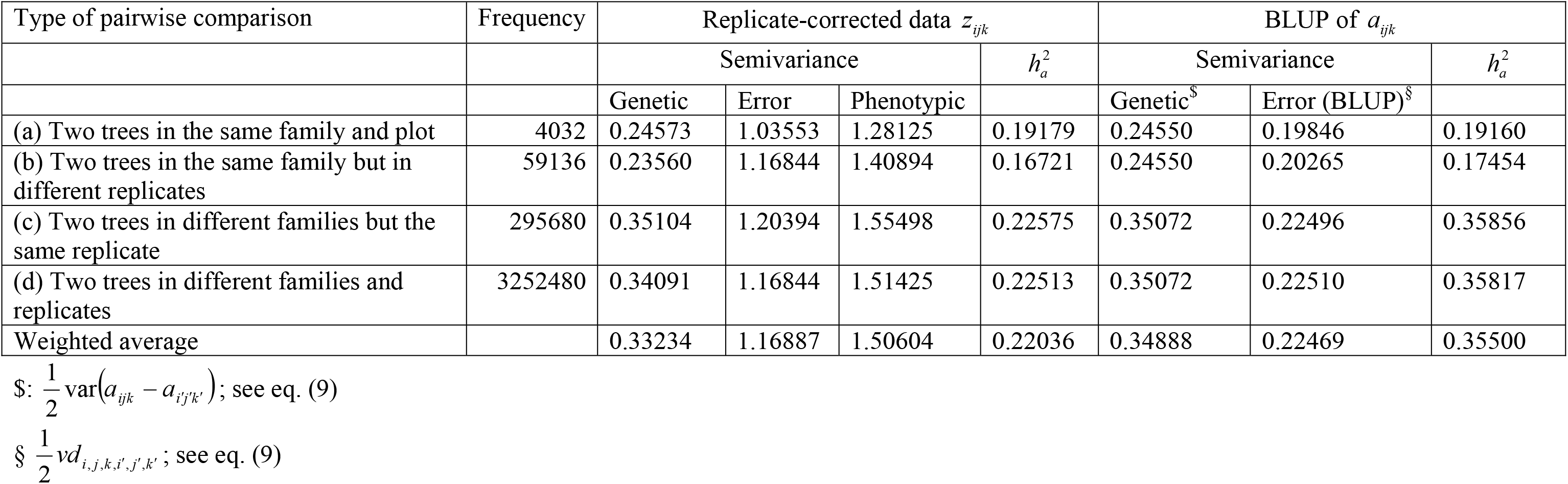
Pairwise heritabilities for four types of comparison (a to d) for dbh1 using simulated complete *A. mangium* data (Example 1) assuming *r* = 0.3 and using models (1) and (2) for RCBD to estimate the variance components and semivariances for both replicate-corrected data *z*_*ijk*_ and BLUP of *a*_*ijk*_.

For comparison, we also computed the mean heritability in eq. (10), in which the averaging is done separately for the numerator and denominator. The result is 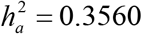, which is close to the average of the pairwise heritabilities based on BLUP in Table 4, which equals 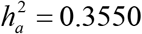, showing that eq. (10) provides a very good approximation.

Figure 1 shows a plot of the BLUPs of genetic effects *a*_*ijk*_ versus the replicate-corrected values *z*_*ijk*_ for dbh1 using simulated complete data. The rank correlation is 0.78. The presence of family effects *f*_*j*_ in the BLUPs of the genetic effects *a*_*ijk*_ = *f* _*j*_ + *b*_*ijk*_ causes a vertical displacement in the same direction of points for the trees in each family, as can be seen by the colour-coding for families in Figure 1.

**Figure 1:**
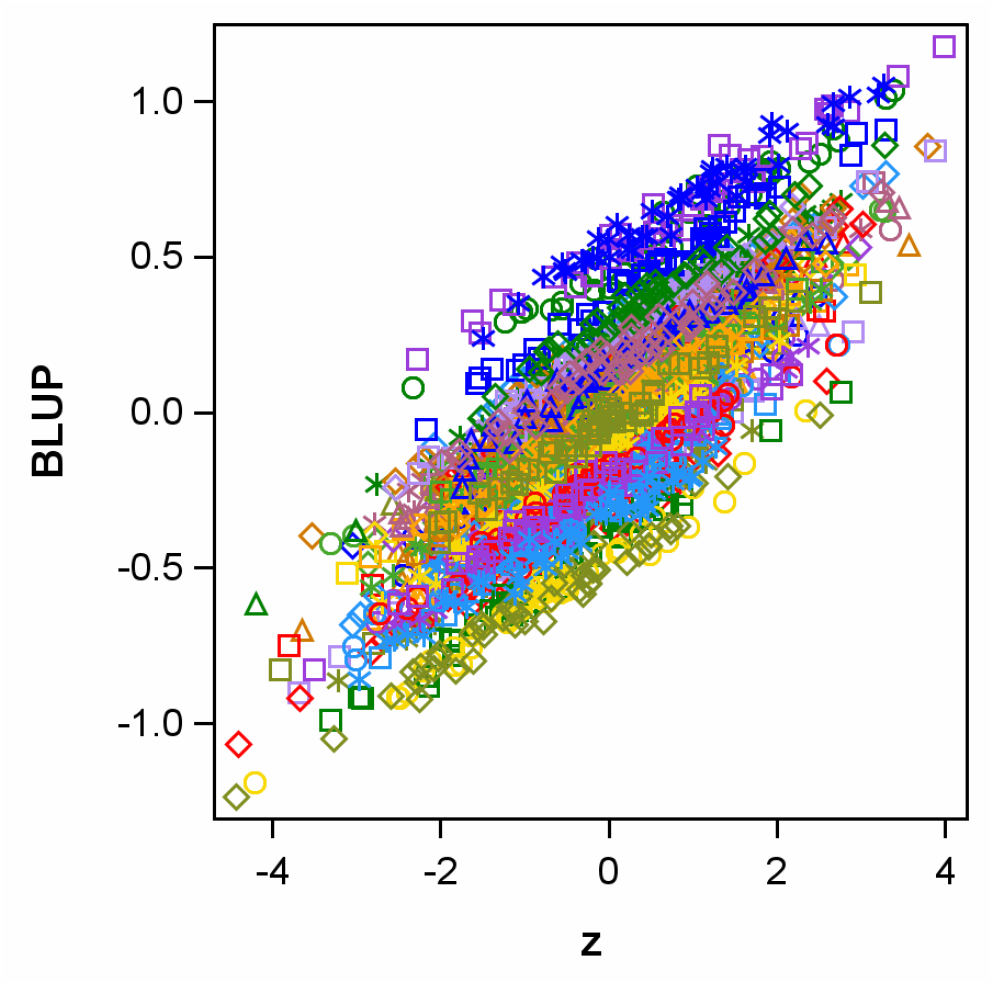
Plot of BLUP of genetic effects *a*_*ijk*_ versus replicate-corrected values *z*_*ijk*_ for dbh1 using simulated complete data. Families are identified by different colours and symbols.

### Incomplete trial data

We also estimated the BLUP-based pairwise heritabilities from the original incomplete trial data for dbh1. The mean of the pairwise heritabilities equals 0.35710, which is similar to the mean heritability for the simulated complete data (Table 4). The histogram of the pairwise heritabilities is shown in Figure 2. There are two clearly separated modes, representing the within-family and between-family comparisons.

**Figure 2:**
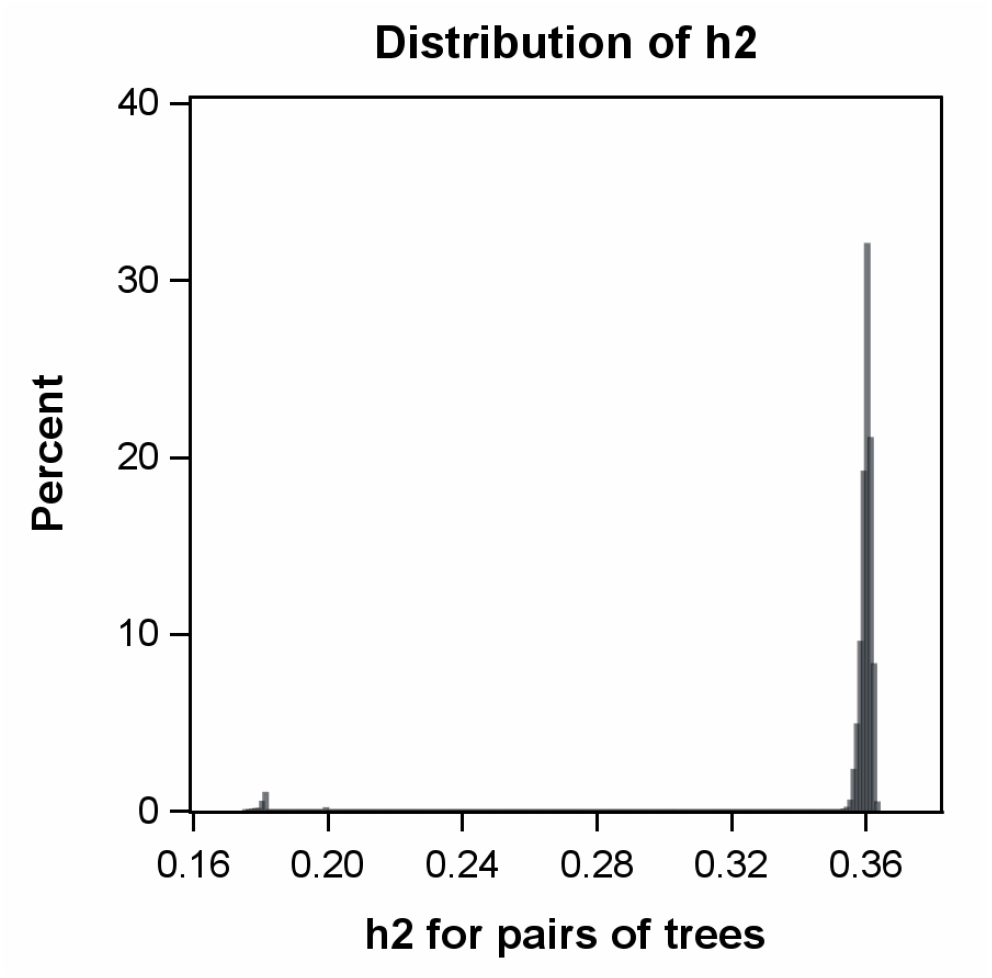
Histogram of BLUP-based pairwise tree-level heritabilities for incomplete dbh1 data based on model (1). The mean of the pairwise heritabilities equals 0.35710.

### Incomplete blocking

Next, consider analysis using the randomization-based model that represents the resolvable row-column design used in the trial. That model includes random row and column effects, nested within replicates. Here, we only report results for the BLUP-based approach using both the incomplete trial data (Figure 3) and the simulated complete data (Figure 4). In both cases, the average heritability based on the randomization-based model is improved compared to the analysis based on model (1) for an RCBD (Figure 2, Table 4). The split in the histograms into within-family and between-family comparisons is similar in all three cases.

**Figure 3:**
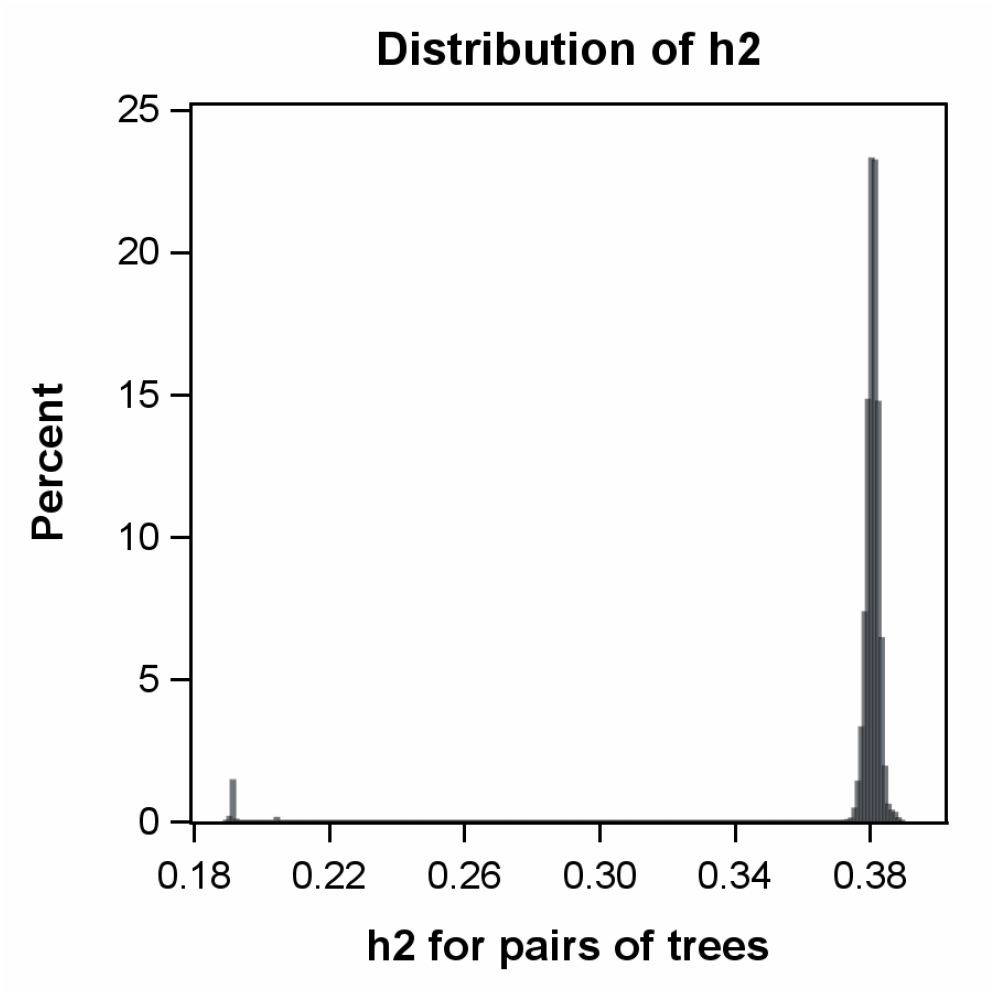
Histogram of BLUP-based pairwise tree-level heritabilities for incomplete dbh1 trial data based on model (1), augmented with random effects for rows and columns, nested within replicates. The mean of the pairwise heritabilities equals 0.37760.

**Figure 4:**
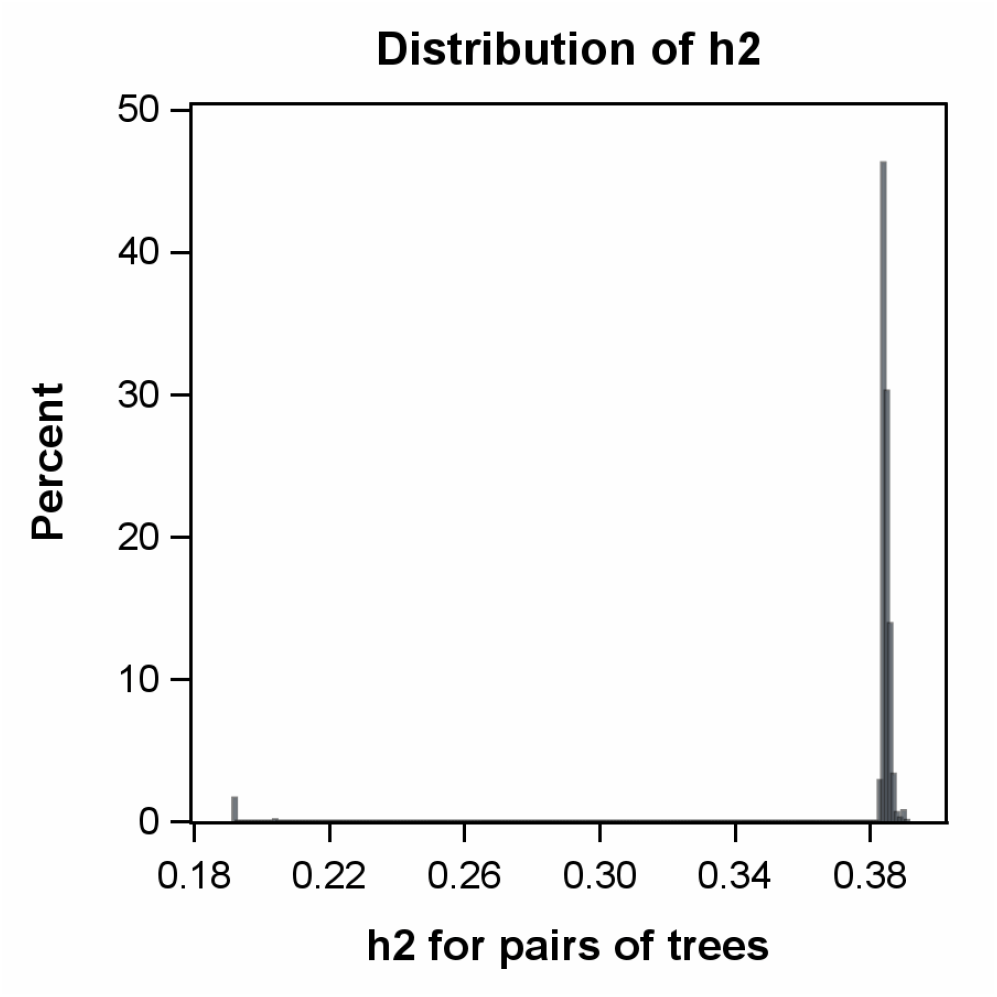
Histogram of BLUP-based pairwise tree-level heritabilities for simulated complete dbh1 data based on model (1), augmented with random effects for rows and columns, nested within replicates. The mean of the pairwise heritabilities equals 0.38138.

## Discussion

In this paper we investigated heritabilities for both the family means and individual trees. For family means, two approaches were considered. The first is based on BLUE of family means and is derived as variance explained by family effects or as regression of family effects on family means. The second is based on BLUP and can be derived as the squared correlation between a random family effect and its estimate. It is noteworthy that for complete data from an RCBD, both definitions lead to the same expression for heritability. For tree-based heritabilities, this equivalence no longer holds. It should be stressed that this does not mean that one of the two approaches is incorrect. The difference just reflects the different philosophies underlying the two approaches, and that a choice needs to be made depending on whether one wants to assess the variance explained by the genetic effect or the main focus is on the reliability of BLUPs. The BLUP-based approach has the advantage that it facilitates the incorporation of any advantages that accrue from incomplete blocks (e.g. rows and columns within replicates).

For individual trees, BLUE is not available. Instead, we considered corrected values obtained by sweeping out replicate effects. Interestingly, heritabilities based on these corrected values, which are also defined in terms of a variance-explained view, were quite different from heritabilities based on BLUP, with clearly higher values for the latter. The only exception is with pairwise heritabilities for two trees in the same plot, where identical results are obtained for two reasons: (i) No correction is needed for block effects and (ii) the pairwise comparison within plots is free of family effects. For comparisons of trees in different families [comparisons (c) and (d) in Table 4], BLUP-based heritabilities are substantially higher for the BLUP-based approach than for the replicate-corrected data. The main reason for this discrepancy is that BLUP can borrow strength across trees in the same family, whereas the replicate-corrected data do not share this property. Inspection of eq. (8) reveals that it is via the BLUP of family effects *f* _*j*_ that information is borrowed from trees in the same family. The heritability associated with this effect,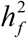, is very high (0.92347 for the simulated complete data), and this explains why BLUP has notably higher average tree-level heritability than the replicate-corrected data. Eq. (8) also explains why comparisons among trees in different families benefit substantially, whereas comparisons among trees in the same family benefit little or not at all from the use of BLUP. Clearly, the BLUP of the differences of two tree-level genetic effects *a*_*ijk*_ is free of *BLUP*(*f*_*j*_), hence no strength is borrowed within families. By comparison, the BLUP of the difference for two trees in different families *j* and *j*′ involves *BLUP*(*f*_*j*_)− *BLUP* (*f*_*j ′*_), so strength is borrowed in this case. We also note that *BLUP*(*f* _*j*_) benefits from efficient blocking. In fact, the experimental designs considered in this paper are optimized for the comparison among family means, which helps the accuracy of *BLUP*(*f*_*j*_)− *BLUP* (*f*_*j′*_) and hence the borrowing of information within families.

We also investigated the gain from incomplete blocking within complete replicates. Our examples based on observed incomplete trial data as well as the simulated data with the same number of trees in each plot demonstrate that there is an improvement in tree-level heritabilities, though the gain is smaller than that from using BLUP in place of replicate-corrected values. Of course, however, the effectiveness of incomplete blocking will depend on the amount of site variation arising from spatial trends in environmental factors such as wind, soil depth, texture and fertility etc. (Williams et al. 2024). It is worth pointing out in this context that our general approach to defining individual tree heritability based on BLUP as set out in the general equations (9) and (10) does not lead to equations that have a simple sum of variance components in the denominator even in the case of complete blocks (Table 2). In case of incomplete blocking, heritability is therefore not expected to be a simple function of the variance component for incomplete blocks. Likewise, in case of spatial modelling, heritability is not a simple function of the residual error variance component. Instead, the efficiency gain from incomplete blocking or spatial analysis is mediated via the relevant portion *C* = var(*â* − *a*) of the inverse of the coefficient matrix of the mixed model equations.

We propose that our general equations (9) and (10) should be used in place of simpler but inappropriate equations for heritability that have been used in the past and that do not allow to build in adjustments that accrue from efficient incomplete blocking.

Model (1) was originally adapted to tree breeding from animal breeding (Borralho 1995; White et al. 2007, Chapter 15), where due to the presence of additive genetic effects *a*_*ijk*_ for individual animals the model is also known as the animal model (Mrode et al. 2023). Our paper has addressed two aspects in relation to this adaptation, which we believe have not been previously considered: (i) the need to incorporate blocking adjustment to cater for within-site variation and experimental design, and (ii) the definition of heritability appropriate for individual tree BLUPs. Regarding blocking adjustment, researchers may additionally consider modelling spatial tree-to-tree variation within and across plots (Dutkowski et al. 2002; Chen et al. 2018). Such model components, which potentially can improve precision, may be used as an add-on to randomization-based analysis, which provides a useful baseline (Piepho and Williams 2010). Furthermore, in relation to the structure of spatial models, there have been some recent spline-based developments as alternatives to autoregressive models (Piepho et al. 2015, 2022). With all of these extensions, the general equations (9) and (10) can be used.

The pairwise heritabilities considered in this paper are primarily useful for descriptive purposes. One might be tempted to use heritabilities proposed in this paper in classical equations to estimate genetic gain (Falconer and Mackay 1996). It needs to be kept in mind, however, that these equations require a balanced design with a symmetric variance covariance structure for what is to be considered the phenotype. For complete data from a RCBD, classical equations to estimate genetic gain can be applied for family means taken as the phenotype. However, for individual trees there is heterogeneity of heritability between pairs of trees, even with a balanced design such as an RCBD with the same number of trees on all plots, as our equations in Tables 1 and 2, as well as the empirical results in Section 3 demonstrate. Hence, classical equations to estimate genetic gain do not apply. A simple way out is to use Monte Carlo simulation, as proposed end exemplified in Piepho and Möhring (2007) and Buntaran et al. (2022). Alternatively, one may average the BLUPs of the selected fraction to assess genetic gain.

## Supplementary Information

The online version contains supplementary material (Appendices S1 and S2) available at …

## Acknowledgements

This work was partially funded by grant PR 1615/3-1 of the German Research Foundation (DFG). We thank Dr Eko Hardiyanto and PT. Musi Huta Persada for providing the *A. mangium* trial data used in Section 3, and Dr Chris Harwood for very helpful discussions and comments on earlier versions of the manuscript.

## Author contribution

The project was conceptualized by Hans-Peter Piepho and Emlyn Williams. Statistical analyses and manuscript writing were conducted by Hans-Peter Piepho and Emlyn Williams. Derivation of semivariances in Table 2 was done jointly by Maryna Prus and Hans-Peter Piepho. Data preparation was conducted by Emlyn Williams.

## Data availability

The *A. mangium* trial data as well as the simulated complete data are available with the Supplementary Information.

