## Appendices S1 and S2 for "Assessing the efficiency and heritability of blocked tree breeding trials"

**Appendix S1: Derivation of BLUP for model (2)**

The variance-covariance matrix for the observed data vector under model (2) can be written as $V=\sigma_{t}^{2}W$ with $W=I_{n}+\gamma_{m}P_{m}+\gamma_{f}P_{f}$, where $I_{n}$ is the *n*-dimensional identity matrix with $n=\text{IJK}$, $P_{m}$ and $P_{f}$ are projection matrices (James and Williams 2024) for plots and families, respectively,

$\gamma_{m}=\frac{{K\sigma}_{m}^{2}}{\sigma_{t}^{2}}=\frac{h_{m}^{2}}{1-h_{m}^{2}}$ with $h_{m}^{2}=\frac{{K\sigma}_{m}^{2}}{{K\sigma}_{m}^{2}+\sigma_{t}^{2}}$, and

$\gamma_{f}=\frac{\text{KI}\sigma_{f}^{2}}{\sigma_{t}^{2}}=\frac{h_{f}^{2}}{\left( 1-h_{m}^{2} \right)\left( 1-h_{f}^{2} \right)}$ with $h_{f}^{2}=\frac{\sigma_{f}^{2}}{\text{KI}\sigma_{f}^{2}+{K\sigma}_{m}^{2}+\sigma_{t}^{2}}$.

Assuming complete data from an RCBD, BLUP of any random effect vector under model (2) can be written as $\text{CV}^{-1}\left( I_{n}-P_{\rho} \right)y$, where $C$ is the covariance among the random effect vector and the data vector $y$ and $P_{\rho}$ is a projection matrix for replicates. The projection matrices $P_{m}$, $P_{\rho}$ and $P_{f}$ will play a key role in subsequent derivations. For example, in our case $P_{m}$ can be written as $P_{m}=I_{I}\otimes I_{J}\otimes K_{K}$ with $K_{K}=\frac{1}{K}1_{K}1_{K}^{T}$. We will make use of the facts that, e.g., $P_{m}P_{m}=P_{m}$, $P_{m}P_{\rho}=P_{\rho}$, $P_{m}P_{f}=P_{f}$ and $P_{\rho}P_{f}=P_{\mu}$, where $P_{\mu}$ is the projection matrix for the grand mean.

To simplify the subsequent derivation, we here assume that the entries in the random effect vector are aligned with those in the data vector $y$. For the family effects this means that each family effect is repeated $\text{KI}$ times on the vector of random effects.

The inverse of $V$ takes the form $V^{-1}=\frac{1}{\sigma_{t}^{2}}W^{-1}$ with $W^{-1}=I_{n}+\theta_{m}P_{m}+\theta_{f}P_{f}$, where

$\theta_{m}=-\frac{\gamma_{m}}{1+\gamma_{m}}=-h_{m}^{2}$ and

$\theta_{f}=-\frac{\gamma_{f}}{\left( 1+\gamma_{m} \right)\left( 1+\gamma_{m}+\gamma_{f} \right)}=-\left( 1-h_{m}^{2} \right)h_{f}^{2}$.

In deriving the BLUPs, we may use $\text{CV}^{-1}=\text{DW}^{-1}$, where $D=\frac{1}{\sigma_{t}^{2}}C$.

(i) Family effects $f_{j}$

$$C_{f}=\text{KI}\sigma_{f}^{2}P_{f}$$

$$D_{f}=\gamma_{f}P_{f}=\frac{h_{f}^{2}}{\left( 1-h_{m}^{2} \right)\left( 1-h_{f}^{2} \right)}P_{f}$$

$\begin{aligned} D_{f}W^{-1}=\frac{h_{f}^{2}}{\left( 1-h_{m}^{2} \right)\left( 1-h_{f}^{2} \right)}P_{f}\left[ I_{n}-h_{m}^{2}P_{m}-\left( 1-h_{m}^{2} \right)h_{f}^{2}P_{f} \right]=\frac{h_{f}^{2}}{\left( 1-h_{m}^{2} \right)\left( 1-h_{f}^{2} \right)}\left[ P_{f}-h_{m}^{2}P_{f}-\left( 1-h_{m}^{2} \right)h_{f}^{2}P_{f} \right]= \\ \frac{h_{f}^{2}}{\left( 1-h_{m}^{2} \right)\left( 1-h_{f}^{2} \right)}\left[ \left( 1-h_{m}^{2} \right)\left( 1-h_{f}^{2} \right) \right]P_{f}=h_{f}^{2}P_{f} \end{aligned}$

$$\text{BLUP}\left( f \right)=D_{f}W^{-1}\left( I-P_{\rho} \right)y=h_{f}^{2}\left( P_{f}-P_{\mu} \right)y$$

where we have used $P_{f}P_{\rho}=P_{\mu}$ with $P_{\mu}$ the projection matrix for the grand mean (James and Williams 2024). In scalar form, we find

$$\text{BLUP}\left( f_{j} \right)=h_{f}^{2}\left( \bar{y}_{i\cdot}-\bar{y}_{\cdot} \right)$$

(ii) Plot effects $m_{\text{ij}}$

$$C_{m}={K\sigma}_{m}^{2}P_{m}$$

$$D_{m}=\gamma_{m}P_{m}=\frac{h_{m}^{2}}{1-h_{m}^{2}}P_{m}$$

$$D_{m}W^{-1}=\frac{h_{m}^{2}}{1-h_{m}^{2}}P_{m}\left[ I_{n}-h_{m}^{2}P_{m}-\left( 1-h_{m}^{2} \right)h_{f}^{2}P_{f} \right]=\frac{h_{m}^{2}}{1-h_{m}^{2}}\left[ \left( 1-h_{m}^{2} \right)P_{m}-\left( 1-h_{m}^{2} \right)h_{f}^{2}P_{f} \right]=h_{m}^{2}\left( P_{m}-h_{f}^{2}P_{f} \right)$$

$$\text{BLUP}\left( m \right)=D_{m}W^{-1}\left( I-P_{\rho} \right)y=h_{m}^{2}\left[ \left( P_{m}-P_{\rho} \right)-h_{f}^{2}\left( P_{f}-P_{\mu} \right) \right]y$$

$$\text{BLUP}\left( m_{\text{ij}} \right)=h_{m}^{2}$$

(iii) Genetic tree effects $b_{\text{ijk}}$

$$C_{b}=\sigma_{b}^{2}I$$

$$D_{b}=h_{b}^{2}I$$

$$D_{b}W^{-1}=h_{b}^{2}\left[ I_{n}-h_{m}^{2}P_{m}-\left( 1-h_{m}^{2} \right)h_{f}^{2}P_{f} \right]$$

$$\begin{aligned} \text{BLUP}\left( b \right)=D_{b}W^{-1}\left( I-P_{\rho} \right)y=h_{b}^{2}\left[ I-h_{m}^{2}P_{m}-\left( 1-h_{m}^{2} \right)h_{f}^{2}P_{f} \right]\left( I-P_{\rho} \right)y \\ h_{b}^{2}\left[ \left( I-P_{\rho} \right)-h_{m}^{2}\left( P_{m}-P_{\rho} \right)-\left( 1-h_{m}^{2} \right)h_{f}^{2}\left( P_{f}-P_{\mu} \right) \right]y \end{aligned}$$

$$\begin{aligned} \text{BLUP}\left( b_{\text{ijk}} \right)=h_{b}^{2} \\ \end{aligned}$$

Taken together, (i), (ii) and (iii) lead to equations (5) to (8) in the main text.

**Appendix S2: Derivation of the error semivariances under model (1)**

Model (1) can be written in matrix form as

$$y=X\delta+\text{Za}+\varepsilon$$

where $\delta\text{\textasciitilde}N\left( 0,G \right)$ is the vector of block fixed effects, $a$ is the vector of breeding values, and

$\varepsilon\text{\textasciitilde}N\left( 0,R \right)$ is the residual vector comprising plot and tree errors. For this model, we have from general results for mixed models

$$\text{var}\left( \hat{a}-a \right)=\left[ Z^{T}R^{-1}Z+G^{-1}-Z^{T}R^{-1}X\left( X^{T}R^{-1}X \right)^{-1}X^{T}R^{-1}Z \right]^{-1}$$

With the simplification $Z=I_{n}$:

$$\text{var}\left( \hat{a}-a \right)=\left[ R^{-1}+G^{-1}-R^{-1}X\left( X^{T}R^{-1}X \right)^{-1}X^{T}R^{-1} \right]^{-1}$$

The variance-covariance matrices $R$ and $G$ take the forms

$R=\sigma_{e}^{2}\left( I_{n}+P_{m}\tilde{\gamma}_{m} \right)\Leftrightarrow R^{-1}=\frac{1}{\sigma_{e}^{2}}\left( I_{n}-\frac{\tilde{\gamma}_{m}}{1+\tilde{\gamma}_{m}}P_{m} \right)=\frac{1}{\sigma_{e}^{2}}\left( I_{n}-\tilde{h}_{m}^{2}P_{m} \right)$ and

$G=\sigma_{b}^{2}\left( I_{n}+P_{f}\tilde{\gamma}_{f} \right)\Leftrightarrow G^{-1}=\frac{1}{\sigma_{b}^{2}}\left( I_{n}-\frac{\tilde{\gamma}_{f}}{1+\tilde{\gamma}_{f}}P_{f} \right)=\frac{1}{\sigma_{b}^{2}}\left( I_{n}-\tilde{h}_{f}^{2}P_{f} \right)$ with

$\tilde{\gamma}_{m}=\frac{{K\sigma}_{m}^{2}}{\sigma_{e}^{2}}$, $\tilde{\gamma}_{f}=\frac{\text{KI}\sigma_{f}^{2}}{\sigma_{b}^{2}}$, $\tilde{h}_{m}^{2}=\frac{{K\sigma}_{m}^{2}}{{K\sigma}_{m}^{2}+\sigma_{e}^{2}}$, and $\tilde{h}_{f}^{2}=\frac{\text{KI}\sigma_{f}^{2}}{\text{KI}\sigma_{f}^{2}+\sigma_{b}^{2}}$.

It can be shown that the following results hold:

$$X^{T}R^{-1}X=\frac{1}{\sigma_{e}^{2}}\left( X^{T}X-\tilde{h}_{m}^{2}X^{T}P_{m}X \right)$$

$$X=I_{I}\otimes1_{J}\otimes1_{K}$$

$$X^{T}X=\text{JK}\times I_{I}$$

$$X^{T}P_{m}X=\text{JK}\times I_{I}$$

$$X^{T}R^{-1}X=\frac{\text{JK}}{\sigma_{e}^{2}}\left( I_{I}-\tilde{h}_{m}^{2}I_{I} \right)=\frac{\text{JK}}{\sigma_{e}^{2}}\left( 1-\tilde{h}_{m}^{2} \right)I_{I}=$$

$$\left( X^{T}R^{-1}X \right)^{-1}=\frac{\sigma_{e}^{2}}{\text{JK}\left( 1-\tilde{h}_{m}^{2} \right)}I_{I}$$

$$X\left( X^{T}R^{-1}X \right)^{-1}X^{T}=\frac{\sigma_{e}^{2}}{\text{JK}\left( 1-\tilde{h}_{m}^{2} \right)}\text{XX}^{T}=\frac{\sigma_{e}^{2}}{\text{JK}\left( 1-\tilde{h}_{m}^{2} \right)}\text{JK}\times P_{\rho}=\frac{\sigma_{e}^{2}}{1-\tilde{h}_{m}^{2}}P_{\rho}$$

$$\begin{aligned} R^{-1}X\left( X^{T}R^{-1}X \right)^{-1}X^{T}R^{-1}=\frac{\sigma_{e}^{2}}{1-\tilde{h}_{m}^{2}}R^{-1}P_{\rho}R^{-1}=\frac{\sigma_{e}^{2}}{1-\tilde{h}_{m}^{2}}\frac{1}{\sigma_{e}^{2}}\left( I-\tilde{h}_{m}^{2}P_{m} \right)P_{\rho}\frac{1}{\sigma_{e}^{2}}\left( I-\tilde{h}_{m}^{2}P_{m} \right)= \\ \frac{1}{\sigma_{e}^{2}\left( 1-\tilde{h}_{m}^{2} \right)}\left( P_{\rho}-\tilde{h}_{m}^{2}P_{\rho} \right)\left( I-\tilde{h}_{m}^{2}P_{m} \right)=\frac{1}{\sigma_{e}^{2}}\left( P_{\rho}-\tilde{h}_{m}^{2}P_{\rho} \right)=\frac{1-\tilde{h}_{m}^{2}}{\sigma_{e}^{2}}P_{\rho} \end{aligned}$$

$$\begin{aligned} \left[ \text{var}\left( \hat{a}-a \right) \right]^{-1}=R^{-1}+G^{-1}-R^{-1}X\left( X^{T}R^{-1}X \right)^{-1}X^{T}R^{-1}=\left( \frac{1}{\sigma_{e}^{2}}+\frac{1}{\sigma_{b}^{2}} \right)I-\frac{\tilde{h}_{m}^{2}}{\sigma_{e}^{2}}P_{m}-\frac{\tilde{h}_{f}^{2}}{\sigma_{b}^{2}}P_{f}-\frac{1-\tilde{h}_{m}^{2}}{\sigma_{e}^{2}}P_{\rho}= \\ c_{1}\left( I_{n}+c_{2}P_{m}+c_{3}P_{f}+c_{4}P_{\rho} \right) \end{aligned}$$

$c_{1}=\frac{1}{\sigma_{e}^{2}}+\frac{1}{\sigma_{b}^{2}}=\frac{\sigma_{t}^{2}}{\sigma_{e}^{2}\sigma_{b}^{2}}=\frac{1}{\sigma_{e}^{2}h_{b}^{2}}$ with $h_{b}^{2}=\frac{\sigma_{b}^{2}}{\sigma_{t}^{2}}$

$$c_{2}=-h_{b}^{2}\tilde{h}_{m}^{2}$$

$$c_{3}=-\frac{\sigma_{e}^{2}h_{b}^{2}\tilde{h}_{f}^{2}}{\sigma_{b}^{2}}=-\left( 1-h_{b}^{2} \right)\tilde{h}_{f}^{2}$$

$$c_{4}=-h_{b}^{2}\left( 1-\tilde{h}_{m}^{2} \right)$$

The inverse, i.e. $\text{var}\left( \hat{a}-a \right)$, will be of the form

$\text{var}\left( \hat{a}-a \right)=\frac{1}{c_{1}}\left( I_{n}+d_{2}P_{m}+d_{3}P_{f}+d_{4}P_{\rho}+d_{5}P_{\mu} \right)$.

As we will be looking at pairwise comparisons, the term in $P_{\mu}$ will generally be dropped out and can therefore be ignored. Hence, we only need to find *d*_2_, *d*_3_ and *d*_4_.

$$\begin{aligned} \left( I_{n}+c_{2}P_{m}+c_{3}P_{f}+c_{4}P_{\rho} \right)\left( I_{n}+d_{2}P_{m}+d_{3}P_{f}+d_{4}P_{\rho} \right)= \\ I_{n}+c_{2}P_{m}+c_{3}P_{f}+c_{4}P_{\rho}+ \\ d_{2}P_{m}+c_{2}d_{2}P_{m}+c_{3}d_{2}P_{f}+c_{4}d_{2}P_{\rho}+ \\ d_{3}P_{f}+c_{2}d_{3}P_{f}+c_{3}d_{3}P_{f}+c_{4}d_{3}P_{\mu}+ \\ d_{4}P_{\rho}+c_{2}d_{4}P_{\rho}+c_{3}d_{4}P_{\mu}+c_{4}d_{4}P_{\rho}= \\ I_{n}+\left( c_{2}+d_{2}+c_{2}d_{2} \right)P_{m}+\left( c_{3}+c_{3}d_{2}+d_{3}+c_{2}d_{3}+c_{3}d_{3} \right)P_{f}+\left( c_{4}+c_{4}d_{2}+d_{4}+c_{2}d_{4}+c_{4}d_{4} \right)P_{\rho}+\left( c_{4}d_{3}+c_{3}d_{4} \right)P_{\mu} \end{aligned}$$

Again, we can ignore the term in $P_{\mu}$*.*

(i) $c_{2}+d_{2}+c_{2}d_{2}=c_{2}+d_{2}\left( 1+c_{2} \right)=0\Rightarrow d_{2}=-\frac{c_{2}}{1+c_{2}}$

(ii) $c_{3}+c_{3}d_{2}+d_{3}+c_{2}d_{3}+c_{3}d_{3}=c_{3}\left( 1+d_{2} \right)+d_{3}\left( 1+c_{2}+c_{3} \right)=0\Rightarrow d_{3}=-\frac{c_{3}\left( 1+d_{2} \right)}{1+c_{2}+c_{3}}$

(iii) $c_{4}+c_{4}d_{2}+d_{4}+c_{2}d_{4}+c_{4}d_{4}=c_{4}\left( 1+d_{2} \right)+d_{4}\left( 1+c_{2}+c_{4} \right)=0\Rightarrow d_{4}=-\frac{c_{4}\left( 1+d_{2} \right)}{1+c_{2}+c_{4}}$

We can now look at the semivariances of pairwise comparisons (a) to (d) as reported in Table 2. In these comparisons, some or all of the projection matrices drop out when computing the semivariances:

(a) $P_{m}$, $P_{f}$, $P_{\rho}$ drop out.

(b) $P_{f}$ drops out.

(c) $P_{\rho}$ drops out.

(d) Nothing drops out.

Observing these facts, the semivariances reported in Table 2 are readily obtained.
